# Timing Matters: Lessons From Perinatal Neurogenesis in the Olfactory Bulb

**DOI:** 10.1101/2024.05.06.592776

**Authors:** Teresa Liberia, Kimberly Han, Sarah Meller, Eduardo Martin-Lopez, Charles A. Greer

**Affiliations:** Department of Neuroscience and Department of Neurosurgery, Yale University School of Medicine, 333 Cedar Street, New Haven, CT 06520

**Keywords:** Olfactory bulb, granule cells, interneurons, neurogenesis, synapse formation, local circuits

## Abstract

In the olfactory bulb odorant receptor specific input converges into glomeruli. Deep to the glomeruli coding of odor information is tuned by local synaptic circuits. Deciphering the dendritic organization of granule cells relative to the secondary dendrites of projection neurons is a pivotal for understanding odor processing. We carried out a detailed interrogation of the granule cells including the timing of neurogenesis, laminar distribution and synaptogenesis between granule cells and projection neurons. In brief, the granule cells develop following a outside in maturation pattern from embryogenesis to adulthood following a developmental continuum. Granule cells born one week after birth exhibit a unique sublayer specific distribution pattern, marking a transition between embryonic or neonatal and adult stages. Integration into reciprocal synaptic circuits occurred 10 days post neurogenesis, We conclude that timing of neurogenesis dictates the anatomical configuration of granule cells within the olfactory bulb, which in turn regulates a preferential synaptic integration with either mitral cell or tufted cell secondary dendrites.

**Summary Statement:** The integration and distribution of granule cells into the olfactory bulb is determined by the timing of neurogenesis. Location of somata shifts from superficial to deep during development.

## INTRODUCTION

The olfactory system, considered the “oldest” of the sensory systems, is critical for essential behaviors including feeding, reproductive activities and recognition of predators, among others (Lyons-Warren et al., 2021). However, the ever-changing olfactory landscape entails a significant challenge to overcome as it requires continuous adaptation for central processing (Sailor et al., 2016, 2017; Wu et al., 2020a; Coppola and Reisert, 2023).

After binding odorant molecules, olfactory sensory neurons send axons that innervate olfactory bulb glomeruli where they synapse with subpopulations of both projection and interneurons whose dendrites arborize within odorant-receptor specific glomeruli (Nagayama et al., 2014; Tufo et al., 2022). The projection neurons, mitral and tufted cells, extend secondary dendrites in a deeper layer of the OB, the external plexiform layer (EPL), where they establish reciprocal dendrodendritic synapses with the apical dendrites of granule cell interneurons (Greer, 1987; Whitman and Greer, 2007; Bartel et al., 2015; Shepherd et al., 2021). The populations of interneurons in the OB are extremely diverse (Figueres-Oñate and López-Mascaraque, 2016) and significantly larger compared to other brain regions. Olfactory interneurons are classified into different classes depending on the location of their soma (for a review see Nagayama et al., 2014). Those located in the granule cell layer (GCL) of the OB, the granule cells (GCs), represent the most abundant (94%) and are a heterogeneous population based on morphology, neurochemistry, connectivity and timing of neurogenesis (Greer, 1987; Woolf et al., 1991; Merkle et al., 2007, 2014; Batista-Brito et al., 2008). The ratio of the GABAergic GCs to excitatory projection neurons is higher than elsewhere in brain; ∼100:1 in the OB (Shepherd et al., 2021), whilst in the neocortex it is ∼5:1 (Markram et al., 2004). GCs are axonless cells with one to few spiny basal dendrites, typically located in the GCL, and an apical dendrite that extensively ramifies into the EPL. Although the absolute number of dendritic spines (gemmules) remains controversial, estimates run from approximately 100 – 300 (counts of spine density are ∼0.3 – 0.4/micron) (Whitman and Greer, 2007; Burton, 2017). Less controversial and based on serial electron microscopy reconstructions is that each granule cell spine establishes a single reciprocal synapse with the secondary dendrite of a projection neuron (Woolf et al., 1991a, 1991b; Woolf and Greer, 1994).

It has been suggested that subpopulations of granule cells may differentially establish synaptic circuits with either tufted or mitral cells. The differential connectivity hypothesis is based on the observation of a segregated ramification pattern of GCs dendritic arbors within the superficial vs. deep portions of the EPL. The dendritic arbor segregation would, in turn, correlate with the location of GC somata in the GCL, superficial-GCL vs deep-GCL, respectively (Mori et al., 1983; Orona et al., 1984; Greer, 1987; Cavarretta et al., 2018).

Despite its complex configuration, the postnatal mammalian OB is highly plastic, primarily due to the life-long influx of neuroblasts coming from the ventricular-subventricular zone (V-SVZ) of the lateral brain ventricles. This quality endows the OB with a mechanism to constantly adapt to a highly variable olfactory environment by providing functionally distinct subpopulations of GCs (Lemasson et al., 2005; Whitman and Greer, 2009; Platel et al., 2019; Tufo et al., 2022).

The V-SVZ, along with the subgranular zone of the dentate gyrus, were first described as neurogenic niches of the adult mammal forebrain in the ‘60’s (Altman, 1962; Altman and Das, 1965; Luskin, 1993; Lois and Alvarez-Buylla, 1994). Work has since focused on understanding the mechanics of this process as well as its impact on circuitry and behavior (Alvarez-Buylla and García-Verdugo, 2002; Sahay et al., 2011; Lepousez et al., 2013; Sakamoto et al., 2014b; Tufo et al., 2022).

Over the past 30 years, extraordinary work has been done to better understand the neurogenic process that supplies the adult OB with new interneurons. As result of this effort, the course of events leading a neuroblast to differentiate into a functional GCs in the OB is reasonably well established. Previous neurodevelopmental investigations from our laboratory demonstrated that adult born GCs requires at least 28 days to achieve mature dendritic growth and peak in spine formation, but only 21 days to begin integrating into reciprocal circuits of the EPL (Whitman and Greer, 2007). Similar studies demonstrated that GCs undergo rapid synaptic integration into the OB circuits as they receive functional excitatory and GABAergic afferents only 3 days after cell birth, significantly prior to the development of output functions (Panzanelli et al., 2009). Survival studies showed that programmed cell death acts as a mechanism that controls adult neurogenesis, suggesting that 50% of all adult-born GCs die between 15 – 45 days post-neurogenesis (Petreanu and Alvarez-Buylla, 2002; Winner et al., 2002; Mechawar et al., 2004), though the topic of programmed cell death among GCs remains controversial (Platel et al., 2019).

Pioneer studies identified the origin and nature of postnatal stem cells in V-SVZ responsible for the continuous production of different types of GCs throughout life (Merkle et al., 2004, 2007, 2014). Other studies, for example, uncovered the relevance of transcription factors, such as Pax6, as intrinsic factors driving neuronal fate (Hack et al., 2005); or the role of functional microglia in the normal development of newborn GCs (Wallace et al., 2020) and their migration into the OB via the rostral migratory stream (Meller et al., 2023). Functionally, interneurons born during adult stages have specific physiological properties that allow them to integrate into the well-established bulbar circuitry following a unique sequence of events (Carleton et al., 2003; Tufo et al., 2022).

Interestingly, GC survival and integration into the olfactory circuits are impacted by sensory experience during a critical period (Carleton et al., 2003; Yamaguchi and Mori, 2005; Katagiri et al., 2011; Livneh and Mizrahi, 2011). In turn, newborn cells play an important role in the formation of olfactory associative memory (Gribaudo et al., 2021) and olfactory learning to discriminate similar odorants (Wu et al., 2020b). Successfully, one mechanism behind this reciprocal influence was uncovered by studies showing that newborn GCs are differentially influenced by olfactory cortical projections during olfactory learning (Lepousez et al., 2014; Wu et al., 2020b). These studies suggest that the continuous addition of new cells to the OB during life, together with the synaptic plasticity that these new neurons experience through early periods during cell maturation, grant the olfactory system the flexibility required to constantly adjust to new behavioral demands (Kelsch et al., 2009).

The effort to understand the mechanisms of adult neurogenesis of OB granule cells has been significant and productive. However, substantially less research has been done on embryonic/early postnatal neurogenesis in mice OB. It should be noted the difference between those two ages, as adult generated neurons must integrate into a completely developed neuronal circuitry, while early generated neurons are born into a developing environment where the neuronal circuits are still developing.

Here, we aim to shed light on OB neurogenesis by first examining the timing of neurogenesis as a determinant mechanism driving the anatomical fate of new GCs. Our results indicate that the differential timing of neurogenesis unequivocally governs the anatomical distribution of GCs in the GCL. Most perinatally generated GCs reside proximal to the internal plexiform layer (IPL) along the superficial-deep axis, while neurons born after full olfactory system development mainly occupy the deepest portion of the GCL. In summary, the GCL develops following a “outside-in” maturation pattern from embryogenesis to adulthood, as a paleocortical structure.

Notably, this developmental pattern has been observed in layer II of the piriform cortex (Martin-Lopez et al., 2019). Moreover, our data revealed that GCs born one week after birth exhibit a unique sublayer-specific distribution pattern, marking a transition between embryonic/neonatal and adult stages. Thus, the addition of GCs to the GCL follows a developmental continuum rather than a dichotomous model. Additionally, we mapped the timeline for synaptic activity of P7-born GCs. Integration into reciprocal circuits occurs 10 days after birth, coinciding with the onset of initial glutamatergic inputs in the apical dendrites of GCs, followed by the establishment of the inhibitory synapses from their apical dendrites to apposed projection neuron secondary dendrites.

## MATERIAL & METHODS

### Animals

All experiments that required BrdU injections were conducted on CD-1 mice (Charles River). For neurogenesis studies, a total of 48 mice were used, males (n=24) and females (n=24). Pregnant females were housed individually, and the day of birth was considered P0. For synaptic integration studies, experiments were conducted on mice produced by cross-mating Ascl1tm1.1(Cre/ERT2)Jejo/J mice (stock #012882, The Jackson Laboratory), maintained as heterozygous (Ascl1^CreERT2/+^), with B6.Cg-Gt(ROSA)26Sortm14(CAG-tdTomato)Hze/J reporter mice (Stock #007914, The Jackson Laboratories). Only littermates with Ascl1-CreERT2/Rosa26-tdTomato genotype were used, which will subsequently be referred as double transgenic mice. In the presence of 4OH-Tamoxifen (4OH-Tx), transit amplifying cells express the fluorescent protein (tdTomato), allowing us to track and study their lineage. Although tdTomato is not a dendritic marker for GCs, it is a reporter protein suited for whole-cell labeling, including cell body and all processes. This experiment randomly included both male and female mice, although lacking any evidence of differences, sex comparisons were not pursued. Mice were housed with a 12-h light/dark cycle with access to standard food and water *ad libitum*. All animal care and use were approved by the Yale University Animal Care and Use Committee.

### 5-Bromo-2’-deoxyuridine (BrdU) and 4OH-Tx injections

To label and analyze the distribution of newly generated interneurons, BrdU (BD Pharmingen) was administrated intraperitoneally (i.p) (50 mg/Kg body weight). A single dose was injected in animals at P0, P7, P21, P55 and P175 at 10 am. To label cells generated embryonically, a pregnant female was injected at gestational day 17 (E17) with one dose i.p. Cells containing BrdU were detected by immunohistochemistry after 50 survival days. The analysis of distribution of BrdU^+^ cells after a survival period of 50 days reveal neuron’s final anatomical location in the GCL.

For analysis of synaptic integration into EPL local circuits, Ascl1-CreERT2/Rosa26-tdTomato mice were used exclusively at P7. These double transgenic animals were separated into 4 groups (n = 3) and injected with a single dose of 4OH-Tx (Sigma-Aldrich). Tissue was collected at 10-, 14-, 17- and 21-days post-injection (DPI) for further analysis.

### Tissue processing

Mice were deeply anesthetized with an overdose of Euthasol (Virbac) and perfused transcardially with 0.9% NaCl in 0.1 M phosphate buffer, pH 7.4 (PBS) with 1 unit/mL heparin, followed by 4% paraformaldehyde (PFA, TJ Baker) in PBS. Animals were decapitated, brains were dissected from the skull and postfixed in the same fixative solution for 1 h at 4 C. Subsequently, they were transferred to PBS overnight and cryoprotected in 30% sucrose in PBS for 48 h. Then, brains were embedded in OCT-compound (Thermo Fisher Scientific). Finally, 25 μm-coronal sections of the OB were serially collected using a Reichert Frigocut Cryostat E-2800. Sections were frozen at −20C until use.

### Immunohistochemistry

Immunohistochemistry was performed on 25-μm-thick coronal sections collected on positive-charged slides. Sections were thawed at 37 C and rehydrated with 0.1 M phosphate-buffered saline (PBS), pH 7.4. Exclusively those containing thymidine analogue were treated with 0.02 N hydrochloric acid (HCl) at 65 C for 35 min for DNA denature. Then, HCL was neutralized by incubating the slides in 0.1 M Boric acid-borax buffer, pH 8.6, at room temperature for 10 min. Only tissue to be stained with gephyrin was treated for antigen unmasking by incubating the slides in 0.01 M citrate buffer, pH 6.0 for 35 min at 65 C. Potential signal from endogenous avidin, biotin and biotin-binding proteins was blocked with Avidin/Biotin Blocking Kit (abcam, ab64212) in those sections where botinylated secondary antibodies were used. Then, all sections were incubated in a blocking solution of PBS supplemented with 0.1% Triton X100 (American Bioanalytical), 5% Normal Goat Serum (NGS, Accurate Chemicals #CL1200-100) or Normal Donkey Serum (NDS, SouthernBiotech #0030-01) and 0.1% Bovine Serum Albumin (BSA, Sigma-Aldrich) for 1 h at room temperature. Next, sections were incubated in primary antibodies diluted in PBS with 0.1% Triton X100 and 1:10 blocking solution. The primary antibodies used are listed in Table 1. After incubation with primary antibodies over night at 4 C, sections were incubated with secondary antibodies (table 1) for 2 h at room temperature in dark conditions. Only those sections incubated with biotinylated secondary antibodies were next incubated with Streptavidin-Alexa Fluor 488 conjugate 1:1000 (ThermoFisher scientific, S32354) in PBS with 0.1% Triton X100. Nuclei were counterstained by incubating the sections with 1 μg/mL of DAPI (Invitrogen) or 5 μM DRAQ5" (BS Pharmingen). After each step, the sections were thoroughly washed in PBS. Finally, the sections were mounted with ProLong Diamond antifade mountant (Thermo Fisher Scientific).

**Table 1.** Primary and secondary antibodies used in the study.

| Anti | Host | Isotype | Dilution | Company |
| --- | --- | --- | --- | --- |
| *Primary Antibodies* |  |  |  |  |
| BrdU | Rat | Monoclonal IgG | 1:300 | Abcam<br>(ab6326) |
| NeuN | Mouse | Polyclonal IgG | 1:500 | Sigma-Aldrich<br>(MAB377) |
| Synaptoporin | Rabbit | Polyclonal IgG | 1:2000 | Synaptic-Systems<br>(102-002) |
| PSD95 | Rabbit | Monoclonal IgG | 1:250 | ThermoFisher<br>(51-6900) |
| Gephyrin | Mouse | Monoclonal IgG1 | 1:300 | Synaptic-Systems<br>(147-021) |
| RFP | Rabbit | Polyclonal IgG | 1:1000 | Rockland<br>(600-401-379S) |

| Anti | Host | Label | Dilution | Company |
| --- | --- | --- | --- | --- |
| *Secondary Antibodies* |  |  |  |  |
| Rat | Goat | A488 | 1:1000 | Invitrogen<br>(A-11006) |
| Mouse | Goat | A555 | 1:1000 | Invitrogen<br>(A-21422) |
| Rabbit | Goat | A488 | 1:1000 | Invitrogen<br>(A-11008) |
| Mouse | Goat | A488 | 1:1000 | Invitrogen<br>(A-11001) |
| Rabbit | Goat | A555 | 1:1000 | Invitrogen<br>(A-21428) |
| Rabbit | Donkey | A555 | 1:1000 | Invitrogen<br>(A-31572) |
| Mouse | Donkey | biotin | 1:1000 | ThermoFisher<br>(A16027) |
| Rabbit | Donkey | biotin | 1:1000 | Jackson<br>ImmunoResearch<br>(715-065-151) |

### Imaging & Quantification

BrdU images were taken with an ORCA-spark Digital CMOS camera (Hamamatsu Model C11440-36U) camera attached to an Olympus BX51 epifluorescence microscope using a 10x objective lens. For each age group a total of 8 animals were imaged (female n = 4, male n = 4) as follows. Along the rostro-caudal axis, 7 to 8 serial sections −250 μm apart - were imaged per animal, from the most rostral section where the GCL and RMS were perfectly identified until the appearance of the anterior olfactory bulb. Using Fiji software, the area comprising MCL, IPL and GCL was divided into 4 equal sublayers of 50 μm width and all the BrdU-containing cells within each sublayer were manually counted using Multi point tool and represented as number of BrdU^+^ cell per mm^2^ (Figure 1. C). Subependymal zone was excluded in all sections analyzed.

**Figure 1.**
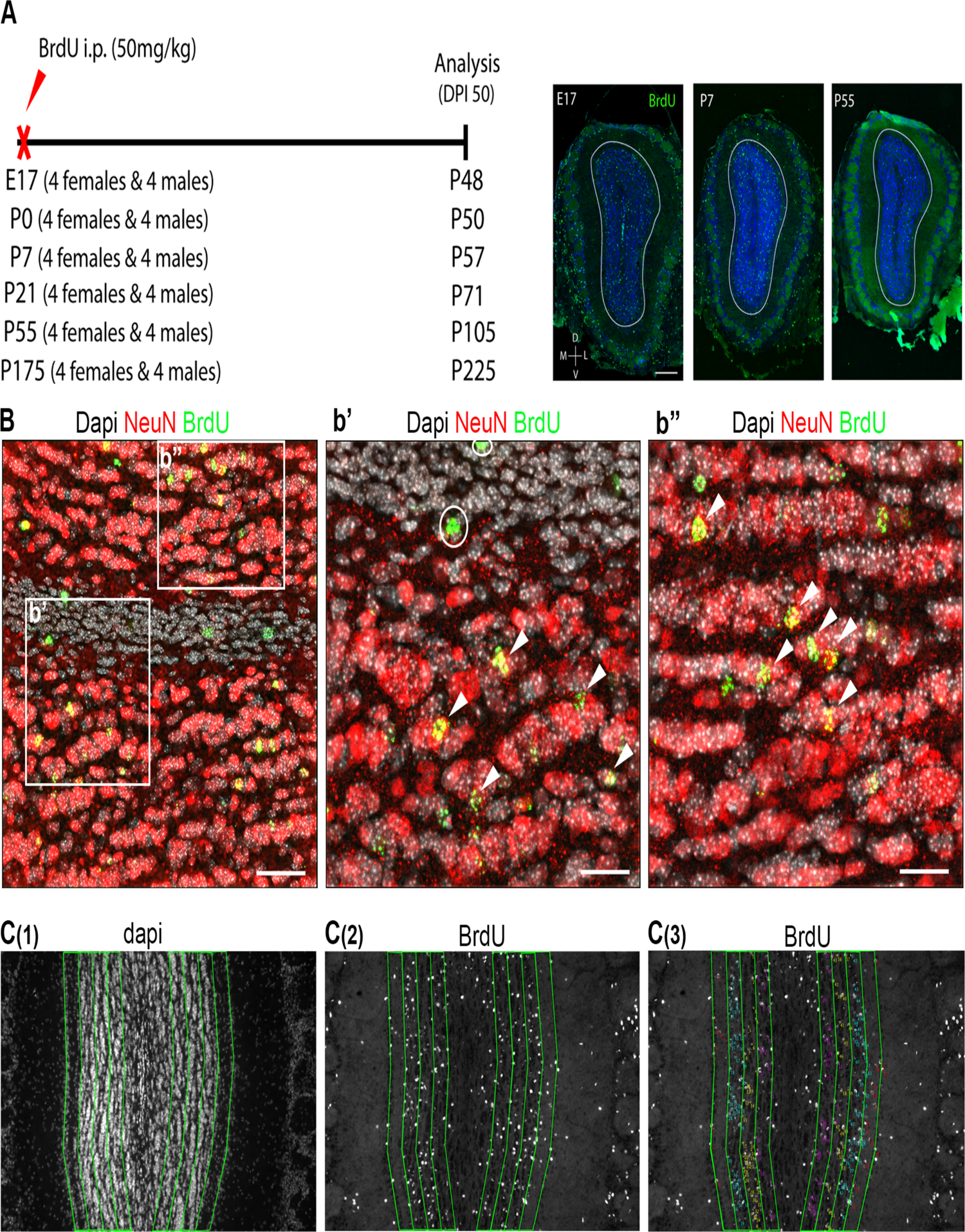
Cellular labeling and quantification strategies of newborn GCs at different embryonic and postnatal time points. (A) Timeline of BrdU injections and analysis. Representative images of coronal sections of the mouse OB show BrdU^+^ cells (green) generated at E17, P7 and P55 respectively. White line delineates MCL. (B) Confocal image shows BrdU^+^ (green) and NeuN^+^ (red) labeling in the GCL and subependymal zone (SEZ). Square areas are magnified in (b’) and (b”) panels. (b’-b”) Double immunofluorescence shows co-expression of BrdU^+^ (green) and NeuN^+^ (red) several GCs in the GCL (arrow heads). Note the lack of BrdU/NeuN co-labelling in the SEZ (circle) in panel (b’). (C) Example of methodology used to analyze BrdU^+^ cell density and distribution across four sublayers encompassing MCL, IPL and GCL in the OB. (C(1)) Coronal OB section counterstained with Dapi (gray) is used to manually delineated a grid consisting of four longitudinal 50-μm-wide sublayers (green lines) covering MCL, IPL and GCL in both medial and lateral sections of the OB. SEZ is excluded. In (C(2)) BrdU^+^ staining (gray) and grid are superposed. (C(3)) Example of Brdu^+^ cell counting method using Fiji software count tool. Scale bar: (A) 300 μm (B) 50 μm. (b’-b”) 25 μm. DPI: day post-injection, D: dorsal, E: embryonic day, L: lateral, M: medial, P: postnatal day, V: ventral.

As it has been previously reported, we recognized different patterns of BrdU labeling during the S phase of the cell cycle. During early S phase, BrdU is associated with dispersed chromatin domains far from the nuclear envelope, revealing a labeling dispersed throughout the nuclear space. However, during late S phase, BrdU labeling is found in perinuclear heterochromatin regions, revealing a ring-like labeling pattern (Mazzotti et al., 1998; Muñoz-Velasco et al., 2013). Both patterns of BrdU labeling were included in our analyses.

Laser scanning microscopy was performed with a Zeiss LSM 800 confocal microscope to analyze the synaptic integration of P7-born GCs into the EPL circuitry. Colocalization between tdTomato-expressing dendrites and three synaptic markers: synaptoporin, gephyrin and PSD-95 were analyzed on 0.2 μm sections collected in z-tacks spanning 2-3 μm in depth using a 63x oil immersion objective. Images were collected from two coronal sections evenly spaced along the anterior-posterior axis for each animal. 3 animals per each time point (10, 14, 17 and 21 days after 4OH-Tx injections) were qualitatively analyzed (Figure 4. A).

### Experimental design and statistical analysis

To study the organization of newborn cells in the GCL, a total of 48 CD-1 mice (24 males and 24 females) were split in six groups of 8 animals (4 males and 4 females). Each group was administered i.p. with a single dose of BrdU (50 mg/kg) at different ages: E17, P0, P7, P21, P55 and P175. After 50 days of survival period, animals were perfused, and BrdU-containing cells were detected by immunohistochemistry.

Multiple sections containing the OB were evaluated per animal as described in “Imaging & quantification” section. The resulting data of the analysis was evaluated to apply the appropriate statistical analysis, which was performed with coded images and blindly to sex and time of BrdU injection. Differences between sexes and among sublayers at and across different time points were assessed by repeated measure (RM) one-way ANOVA or two-way ANOVA (with Tukey’s or Bonferroni’s post hoc test), when appropriate. For a more realistic comparison of number of BrdU^+^ cells in each sublayer across different time points, the results are presented as a percentage (Figure 2. H). Individual tests for each experiment are specified in detail in the results section and table 2. All the statistical analyses were performed using Prism 9 (GraphPad Software). All measures across different sections within an animal were averaged to generate one sample replicate for statistical analysis. For all the statistics, the number of animals in each group was considered as “n”. Data are shown as mean ± SEM. The significance level was set as p-value < 0.05.

**Figure 2.**
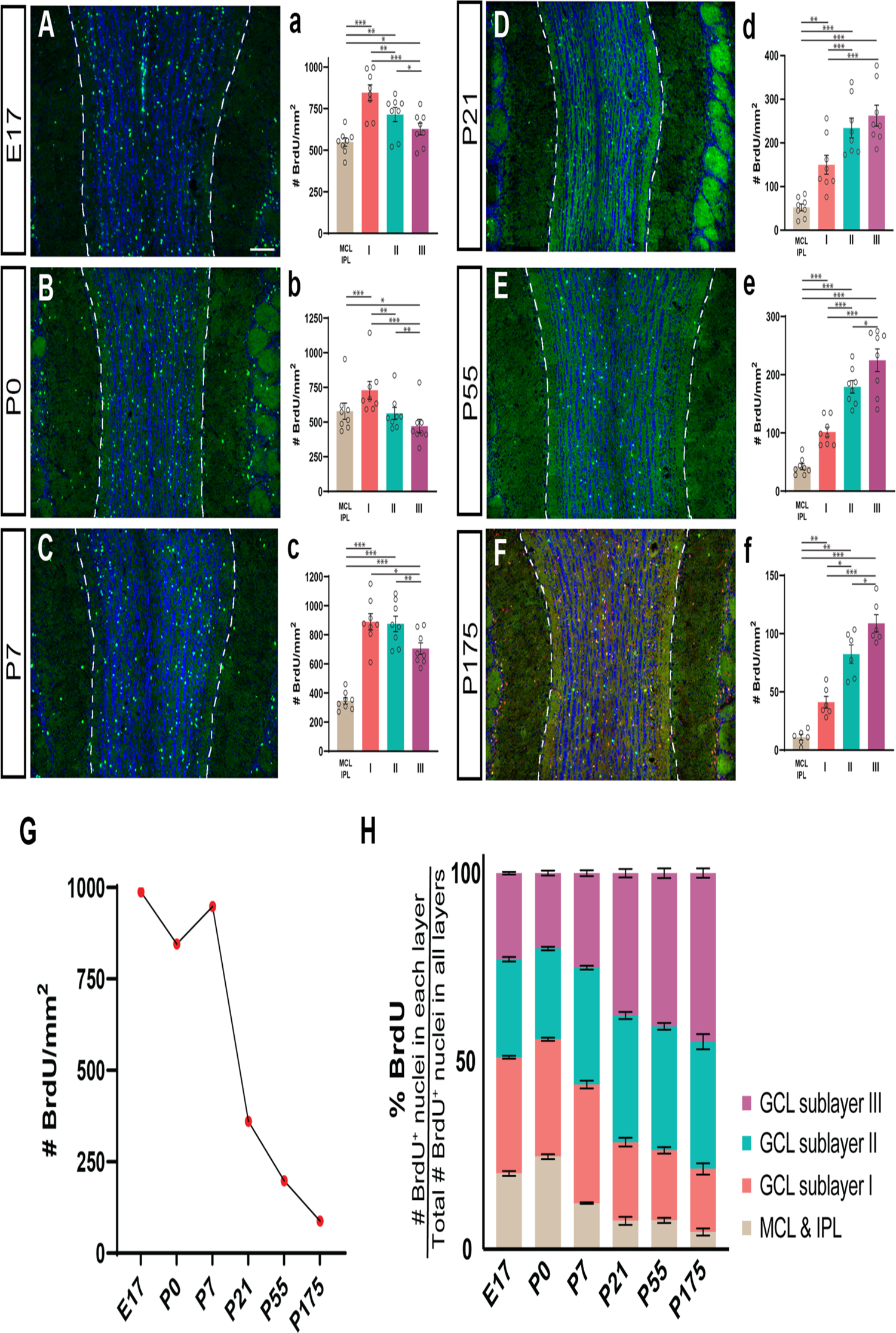
Granule cells generated at different developmental and postnatal stages show different distribution patterns across MCL, IPL and GCL. (A-F) OB coronal sections from CD1 mice showing the spatial organization of granule cells born at E17, P0, P7, P21, P55 and P175 (BrdU^+^). Dotted line demarcates the MCL. (a-f) Graphics illustrating the number of BrdU^+^ cells in MCL & IPL, sublayer I, II and III at different time points. Delineation of sublayers and cell quantification is described in material and methods section. The mean ± SEM values are plotted (n = 8, per each time point) *p < 0.05, **p < 0.01, ***p < 0.001. (G) Graph showing a decrease in granule cells neurogenesis (represented as total number of BrdU^+^) over time. (H) Sublayer-specific distribution pattern of BrdU^+^ granule cells at six different developmental ages. The percentage of newborn granule cells (BrdU^+^) is defined by the ratio between the number of BrdU^+^ nuclei in each sublayer and the total number of BrdU^+^ cells in all sublayers for each age. Scale bar: (A-F) 100 μm. E: embryonic day, GCL: granular cell layer, I: GCL sublayer I, II: GCL sublayer II, III: GCL sublayer III, IPL: internal plexiform layer, MCL: mitral cell layer. P: postnatal day.

**Table 2.**
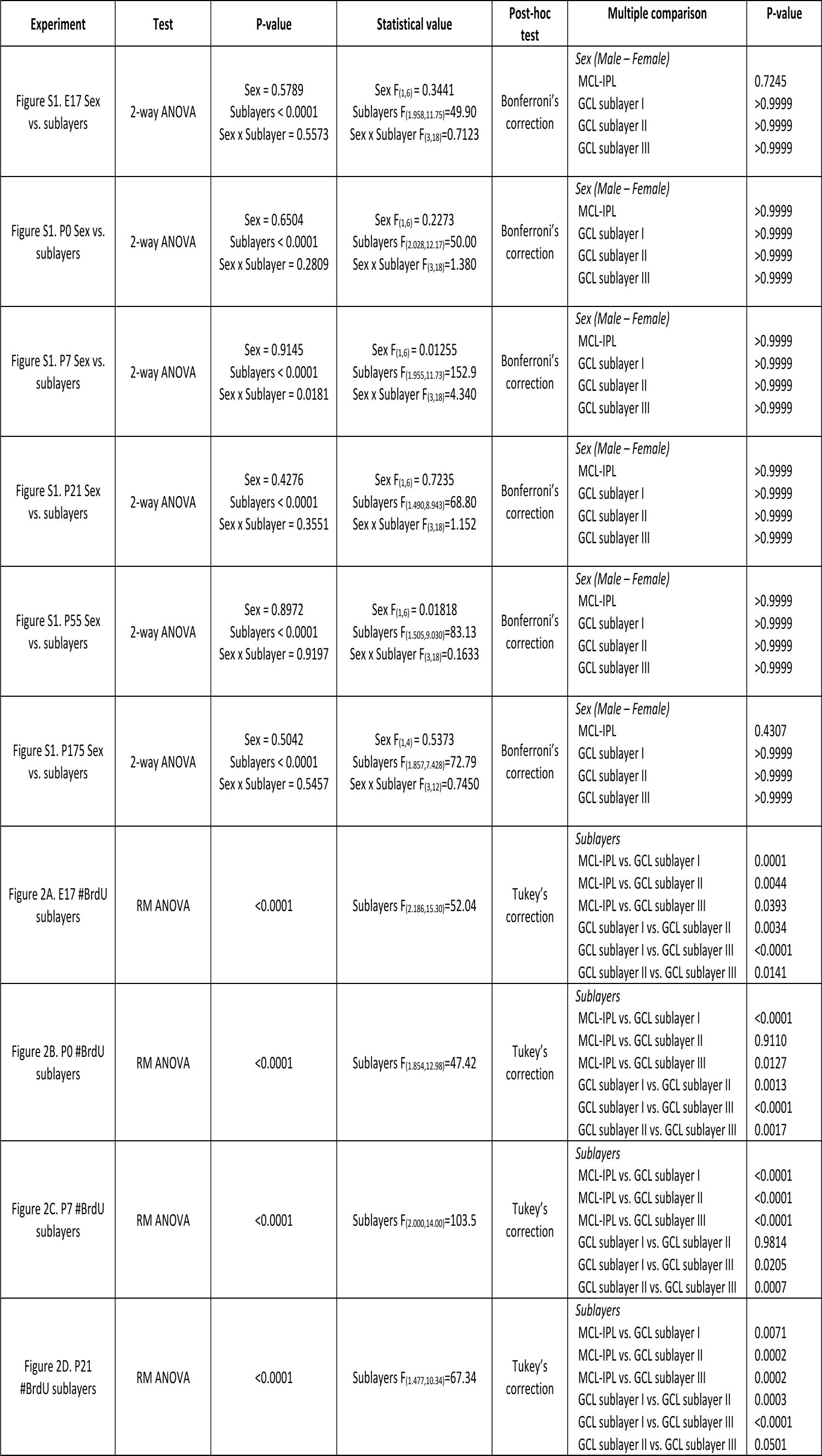
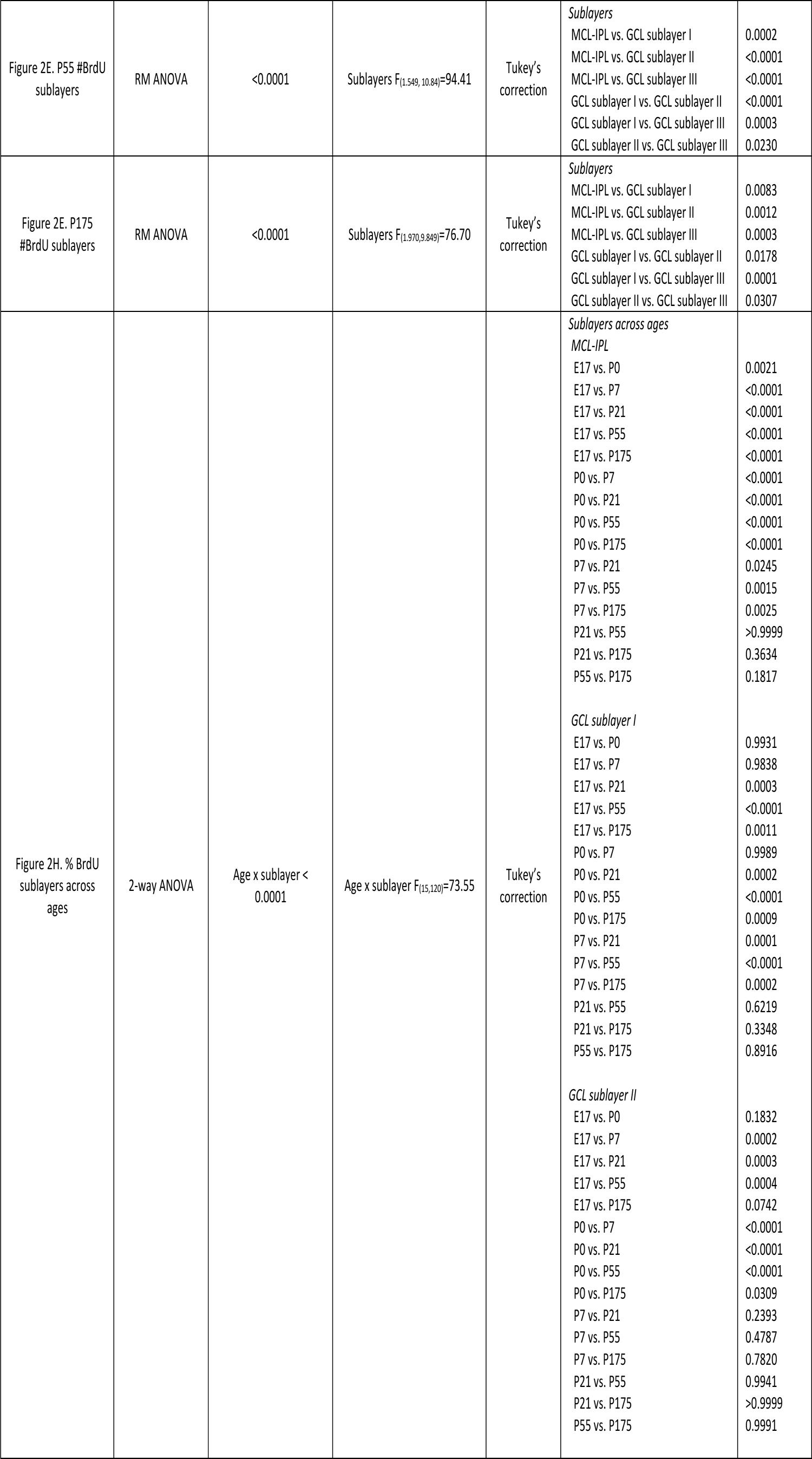
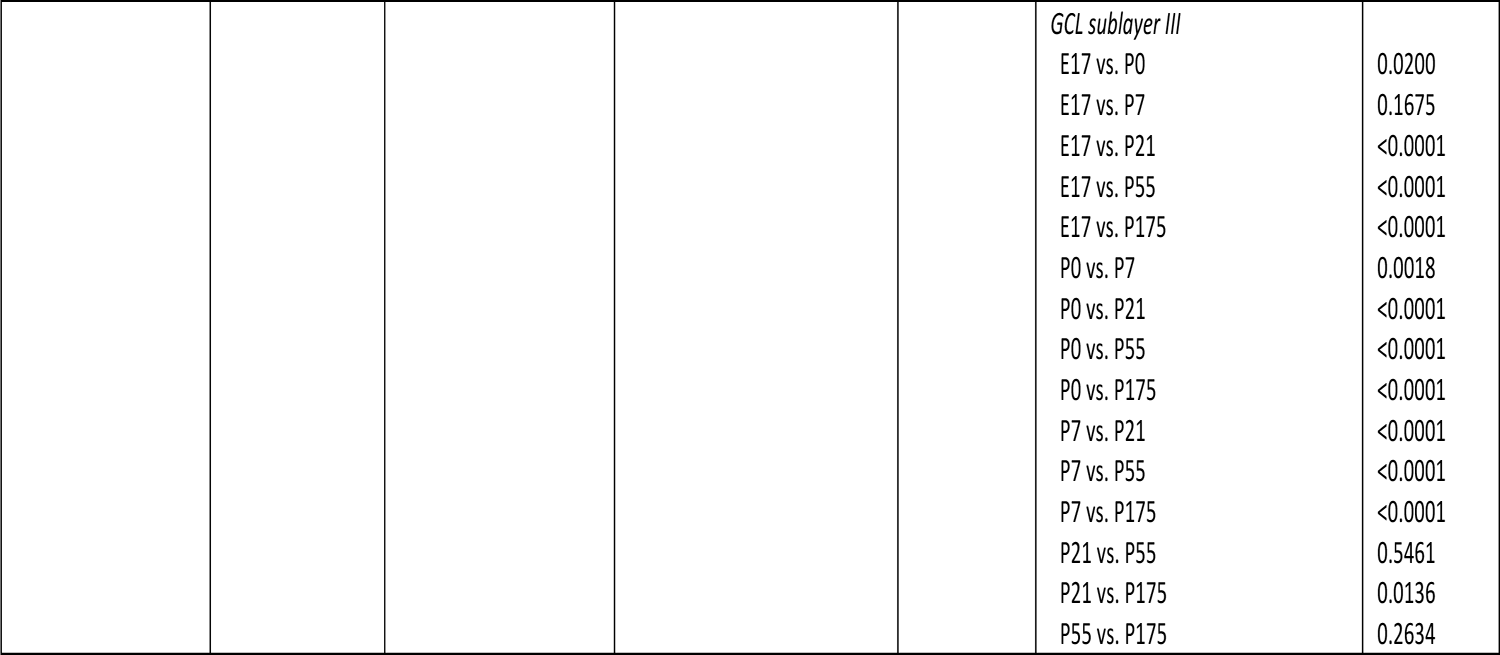
Summary of statistical analysis performed in each experiment. **Abbreviations.** #: number, E: embryonic day, GC: granule cell, GCL: granular cell layer, IPL: internal plexiform layer, MCL: mitral cell layer. P: postnatal day, RM: repeated measures.

## RESULTS

### GC fate based on timing of neurogenesis

To investigate timing of neurogenesis as a potential determinant of GCs fate, we labeled actively dividing cells in the developing and adult brain. To accomplish this, we intraperitoneally (i.p.) injected mice with one dose of BrdU (50 mg/kg of body weight) at one of these developmental time points: embryonic day (E) 17, postnatal day (P) 0, 7, 21, 55 and 175.

Although GCs populate the GCL extensively, they also densely populate the mitral cell layer (MCL) densely (Parrish-Aungst et al., 2007). Therefore, we analyzed their spatial distribution in a broad area comprising the MCL, the IPL and the GCL 50 days post-injection (DPI). Both sexes were examined and compared for each time point.

In the mouse brain olfactory neurogenesis occurs during neurodevelopment and extends throughout the life span, taking place in different regions depending on the age at which it occurs. Olfactory interneurons are produced in the lateral ganglionic eminence and subventricular zone of the lateral ventricle during embryonic and postanal stages, respectively. In both cases, neuroblasts travel long distances to finally reach their destination in the OB. Postnatally, neuroblasts tangentially migrate throughout the rostral migratory stream (RMS) towards the OB. After reaching the subependymal zone (SEZ), they shift to radial migration which allows them to detach from the RMS and reach the GL or GCL, where they will integrate into the local neuronal circuits. The focus of this study is to analyze the distribution of mature GCs throughout the MCL, IPL and GCL. The vast majority of postally generated interneurons require 4 weeks to reach the GCL (Lemasson et al., 2005; Whitman and Greer, 2007). However, Petreanu and Alvarez-Buylla, (2002) estimated that 50% of them die shortly after maturing (15 – 45 days), although this remains controversial (Platel et al., 2019).

Here, to ensure that only mature GCs integrated in the olfactory circuitry are included in this study, we analyze newly born GCs 50 days after BrdU injections. We ensured that they already differentiated into GCs by co-staining for BrdU and NeuN, a neuronal marker expressed by mature GCs and PGs in the OB, but not in M/T (Mullen et al., 1992). First, we verified that BrdU^+^ cells co-expressed NeuN, including those located in the most inner portion of the GCL (Figure. 1 B, b’ and b” arrowhead). Therefore, we considered that BrdU^+^ cells in the GCL, IPL and MCL are differentiated into GCs, contrary to other BrdU^+^ cells located in the SEZ, which were NeuN^-^ (Figure. 1 B, b’ circle), indicating that they did not differentiate into neurons. Our results showed that the number of new generated GCs reaching their final destination in the GCL is negatively correlated with age (Figure 2 G). This is consistent with previous data demonstrating that the rate of GCs neurogenesis decreases over time (Hinds, 1968; Lemasson et al., 2005).

When we examined the distribution pattern of BrdU^+^ GCs in the superficial to deep axis, we observed that it was significantly different depending on their timing of neurogenesis. For each time point, we analyzed the location of BrdU^+^ cells distributed along four equal sublayers encompassing from MCL to GCL (Figure 1 C) in the superficial-deep axis. Each one of these 4 sublayer was identified as: MCL/IPL, GCL sublayer I, GCL sublayer II and GCL sublayer III, respectively, from the most superficial division to the deepest. SEZ was excluded. Statistically significant differences in the number of BrdU^+^ cells populating each sublayer were observed at each time point analyzed (Figure 2 a-f and Table 2). However, no differences between sex were found when the number of BrdU^+^ cells were analyzed at each timepoint (Supplemental Figure S1). At E17 and P0, statistically significant differences were noticed among all the sublayers; with the largest number of BrdU^+^ cells located in GCL sublayer I (Figure 2 a and b, Table 2).

However, a different distribution pattern was observed when GCs born at P7 were analyzed. Here, most newborn interneurons were targeted to GCL sublayer I and II indistinctively (Figure 2 c, Table 2). Finally, a clearly different configuration was found when we analyzed the distribution of GCs generated at P21, P55 or P175. Significant differences in the distribution of BrdU^+^ cells among all the sublayers were seen at each adult time point, being the GCL sublayer III the most populated in all of them (Figure 2 d, e and f, Table 2).

Note that the absolute number of BrdU^+^ cells dropped significantly over time (Figure 2G). Then, to further, and more realistically, compare the sublayer-specific distribution pattern of BrdU^+^ cells across six time points of neurogenesis, we calculated the ratio between the number of BrdU^+^ nuclei in each sublayer and the total number of BrdU^+^ cells in all layers for each time point. Interestingly, statistical analysis showed that the distribution pattern of GCs generated at P7 shared common traits and showed significant differences when compared to interneurons born at P0 and P21 in a sublayer specific fashion. Statistically significant differences were observed in MCL-IPL, CCL sublayer II and III when the ratio of newborn GCs was compared between P7 and P0, but no significant difference was observed in the GCL sublayer I (P0: 25% of BrdU^+^ cells were found in the MCL/IPL, 31% in the GCL I, 24% in the GCL II and 20% in the GCL III. P7: 12% of total BrdU^+^ cells were found in the MCL/IPL, 32% in the GCL I, 31% in the GCL II and 25% in the GCL III) (Figure 2 H, Table 2). In contrast, when percentages of BrdU^+^ cells were compared between P7 and P21, significant differences were observed in MCL-IPL, GCL sublayer I and III, but not in GCL sublayer II (P21: 8% of BrdU^+^ cells were located in the MCL/IPL, 21% in the GCL I, 34% in the GCL II and 37% in the GCL III) (Figure 2 H, Table 2).

Statistical analysis showed that the sublayer-specific distribution pattern was not significantly different when embryonic (E-17) and neonatal time points (P0) were compared (E17: 20% of BrdU^+^ cells were located in the MCL/IPL, 31% in the GCL I, 26% in the GCL II and 23% in the GCL III) (Figure 2 H, Table 2). Equivalent not statistically significant results were observed when the population of GCs born at P21, P55 and P175 were compared (P55: 8% of BrdU^+^ cells were found in the MCL/IPL, 19% in the GCL I, 33% in the GCL II and 40% in the GCL III. P175: 5% of BrdU^+^ cells were found in the MCL/IPL, 17% in the GCL I, 34% in the GCL II and 44% in the GCL III) (Figure 2 H, Table 2).

In summary, our results showed that the ongoing influx of GCs contributes to the development of the OB along a spatiotemporal continuum. Our data suggest P7 as a key transition point, as the distribution of P7-born GCs along the superficial-deep axis represents a transition between the organization pattern seen at earlier developmental times vs. adult stages.

### Integration of P7-generated GCs into EPL circuitry

To consider newborn GCs fully mature they need to meet different criteria. They are required to: (1) reach their final destination in the OB; (2) extend their dendritic arbors profusely in the EPL while developing their numerous dendritic gemmules; and (3) integrate into the synaptic circuitry by stablishing reciprocal dendro-dendritic synapses with bulbar projection neurons (MCs and TCs). In the EPL, MCs restrict their secondary dendrites within the deepest portion of the EPL, while TCs ramify theirs along the outer EPL (Mori et al., 1983; Orona et al., 1984). In a similar fashion, GCs dendritic arbors seem to be segregated in superficial or deep portions of the EPL based on the location of their soma in the GCL (Greer, 1987). Previous studies have shown that MCs and TCs are affected differently by lateral inhibition as they differentially recruit morphological distinct classes of GCs (Geramita et al., 2016). The anatomical segregation of dendrites of projection neurons and GCs in the EPL suggests that GCs differentially connect with TCs or MCs based on their location in the superficial or deep GCL. This is consistent with the idea that the olfactory system uses parallel pathways to encode different features of the olfactory information.

In this study we showed that the final anatomical location of GCs along the superficial-deep axis is determined by timing of neurogenesis. Most notably, we demonstrated that P7-newborn GCs showed a unique distribution pattern in the OB, which was significantly different from the distribution of interneurons produced embryonically, neonatally, and from those generated later, when the olfactory system is fully developed. To further understand the behavior of this subset of interneurons, we investigated their timeline of synaptogenesis as a proxy for their functional integration into the brain circuits.

Although, previous work has shown that early postnatally-generated interneurons integrate into olfactory circuits following a different pattern than adult-born GCs in the rat OB (Kelsch et al., 2008), no data about the synaptic integration timeline of perinatally generated interneurons has been collected in mouse OB. We cannot assume that neurogenic processes are similar between rats and mice, as Snyder and colleagues demonstrated in previous studies of hippocampal neurogenesis (Snyder et al., 2009). Here we attempted to fill this gap in our understanding of the developmental determinants and integration of perinatally produced granule cells.

To study the synaptogenesis timeline of P7-born GCs, we first labeled them by using *in vivo* lineage tracing with inducible Cre recombinase. In the postnatal brain, Ascl1, a bHLH transcription factor, is expressed in transit amplifying cells that give rise to GABAergic interneurons (Kim et al., 2007) (cf. Supplmental Figure 2). This strategy allows us to: (1) label interneurons generated specifically at the time of 4OH-Tx injection: and (2) label them in their entirely because the fluorescent protein tdTomato is expressed in every compartment of a cell: soma, axon, dendrites and dendritic specializations. Then, we analyzed the onset of synaptic contacts on GCs tdTomato^+^ dendrites. To accomplish that, we immunostained for synaptic associated proteins: synaptoporin, gephyrin and PSD-95 (Figure 3. A and B).

**Figure 3.**
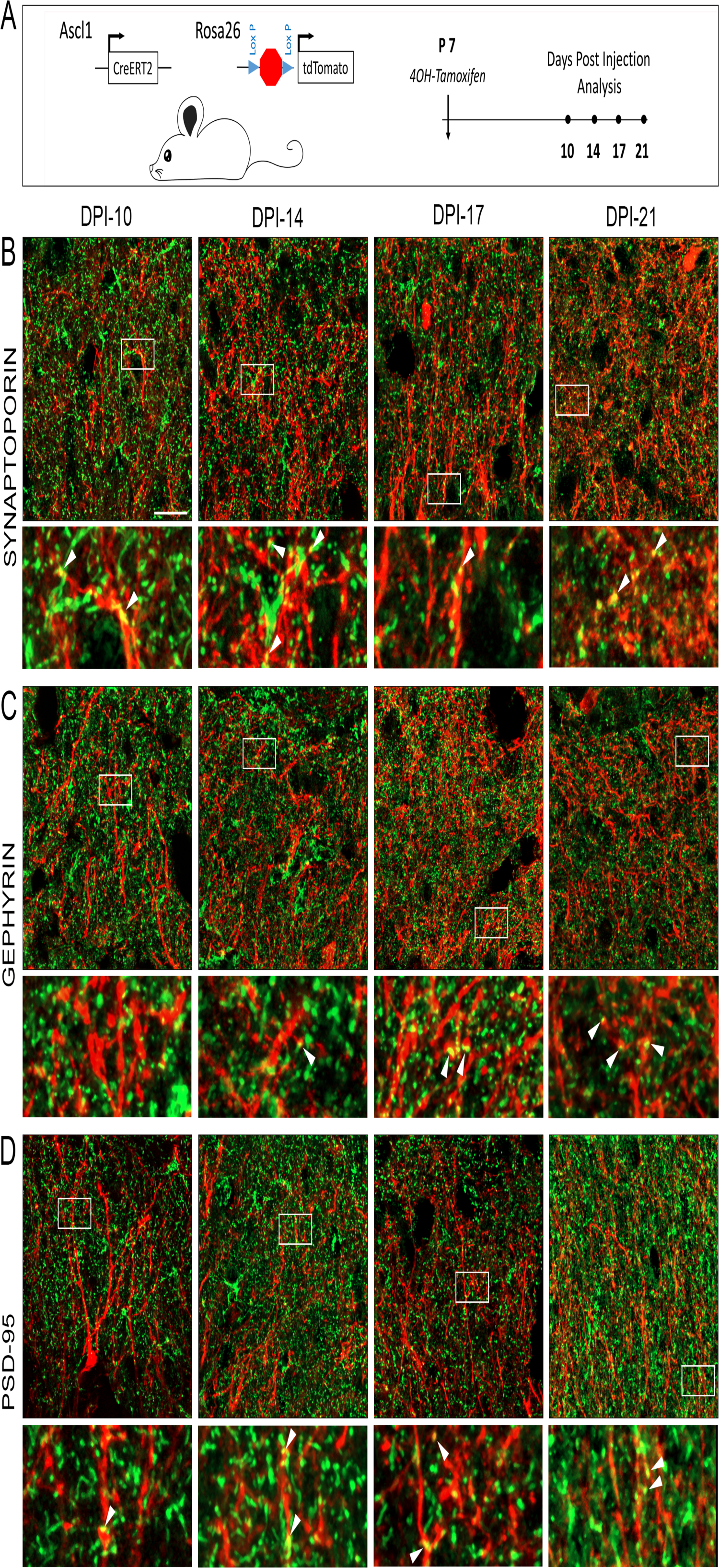
In vivo genetic fate-mapping strategy reveals spatiotemporal synaptic integration of P7-born GCs. (A) Experimental paradigm using Inducible Cre-lox P strategy to label GCs generated at P7 in Ascl1-CreERT2/Rosa26-tdTomato double transgenic mice and analysis timeline. (B-D) High-magnification confocal images representing flattened stacks of two to three 0.2-μm-thick optical sections showing tdTomato^+^ GCs dendrites (red) within the EPL at DPI-10, −14, −17 and −21. Immunohistochemistry for presynaptic and postsynaptic markers (green): (B) synaptoporin, (C) gephyrin and (D) PSD-95 at different timepoints after 4OH-Tx injection shows the chronology of synaptic integration of P7-born GCs into EPL local circuits. (B) tdTomato^+^ dendrites colocalize with synaptoporin synaptic marker at DPI-10 for the first time. (C) Gephyrin^+^ puncta appear juxtaposed to tdTomato^+^ dendrites consistently as soon as DPI-14, while (D) PSD-95^+^ puncta colocalize with tdTomato^+^ dendritic spines at DPI-10. In (B,C and D), white rectangles indicate the area shown at higher magnification below each panel for each marker and timepoint. White arrowheads indicate examples of double-label. Scale bar: 20 μm. DPI: day post injection, EPL: external plexiform layer.

As GCs lack axons, they are reliant on their dendritic arbor for their output function. Their dendrites make dendrodendritic synaptic contacts with MCs and TCs within the EPL as their principal mode of connectivity. In many cases, those dendrodendritic contacts are reciprocal (Greer, 1987; Shepherd et al., 2021). This means that the presynaptic element in an excitatory synapse is, at the same time, the postsynaptic element in an inhibitory synapse, and vice versa. Moreover, GCs receive axosomatic and axodendritic inputs to their cell bodies and basal dendrites, respectively, from MCs/TCs axon collaterals and centrifugal fibers from the olfactory cortex and modulatory structures (Shepherd et al., 2021).

In the present study we focused on the development of synapses in the EPL where the majority of synaptic activity involving GCs occurs. First, we analyzed the expression of synaptoporin, a marker for mature granule cell spines, at different time points after 4OH-Tx injection as a readout of when newborn GCs engage in the olfactory synaptic activity for the first time. Synaptoporin is a membrane protein of synaptic vesicles localizing specifically in dendritic spines of GCs, but not in MCs (Bergmann et al., 1993; Whitman and Greer, 2007b; Breton-Provencher et al., 2016). We found the presence of tdTomato^+^ dendritic spines expressing synaptoporin at 10 dpi, as well as at 14, 17 and 21 dpi, where the presence of colocalization becomes more evident (Figure 3. B). Note that puncta-like synaptoporin staining pattern in the EPL is consistent with the elevated synaptic activity occurring in the EPL.

To further analyze whether P7-generated GCs are able to form inhibitory synaptic contacts from their dendritic spines and to determine when it occurs, we analyzed the expression of gephyrin. Gephyrin is a key protein that anchors and clusters glycine and γ-aminobutyric acid type A (GABA) receptors at inhibitory synapses. It also plays an important role in neuronal growth and synapse formation (Sassoè-Pognetto and Fritschy, 2000; Deprez et al., 2015). As GCs make inhibitory contacts with dendrites of MCs/TCs, the expression of this postsynaptic marker is expected to be found in MCs/TCs dendrites, and thus, in juxtaposition with tdTomato^+^ dendritic spines. We found gephyrin^+^ puncta in close apposition to labeled GCs spines 14 days after 4OH-Tx injection, which became more obvious in later time points, 17 and 21 dpi (Figure 3. C). However, gephyrin^+^ puncta juxtaposed with labeled spines at 10 dpi were extremely rare (Figure 3. C).

As noted, the dendritic compartment of GCs represents the postsynaptic element in asymmetrical synaptic contacts with projection neurons. A significant percentage of excitatory synapses in the brain occurs exclusively in dendritic spines (Nimchinsky et al., 2002). However, olfactory GCs represent again an exception to the norm and some glutamatergic synapses are also found on their dendritic shaft (Woolf et al., 1991b; Sailor et al., 2016; Shepherd et al., 2021). To further characterize the synaptogenesis time course of P7-generated GCs, we analyzed when they receive excitatory inputs in the EPL. Since MCs/TCs release glutamate at the dendritic synapse sites, we use PSD 95 as a postsynaptic marker to study excitatory inputs in labeled GC dendrites. We found PSD95^+^ puncta in the EPL colocalizing with tdTomato^+^ dendrites as soon as 10 days after 4OH-Tx injection (Figure 3. D).

In rats, adult-born GCs require more time to integrate into the EPL local circuits when compared to neonatally-born GCs. Likewise, the synaptic distribution pattern differs depending on when the interneurons are born. Adult-born GCs receive glutamatergic inputs in their proximal dendrite several days before they do in the distal portion of the apical dendrite. However, neonatal GCs receive synaptic inputs simultaneously in their apical dendrites in the EPL and in their basal dendrites in the GCL (Kelsch et al., 2008). In the mouse, adult-born GCs require at least 21 days to first integrate into the EPL circuitry (Whitman and Greer, 2007). Our results showed that, in the mouse OB, early developmentally-born GCs (P7) integrated into the EPL circuitry much faster than the previously reported newborn cells produced during adult periods (Whitman and Greer, 2007).

Morphologically, we observed that P7-generated GCs extended and ramified their apical dendrites into the EPL over time. At DPI-10 relatively unbranched apical dendrites were detected in the EPL (Figure 4. A). Those dendrites branched and ramified into the EPL by DPI-17 and thereafter, when they exhibited a more evident spiny appearance (Figure 4. B-C). It is important to highlight that spiny basal dendrites were not obvious in perinatally born GCs until DPI-17 (Figure 4. G-I). Likewise, long neck spines in apical dendrites became more apparent 17 days after cell birth and beyond, remaining sparse at DPI-10 (Figure 4. D-F). This is consistent with the timeline of synaptic vesicle-associated protein expression, where PSD-95 expression was first detected in apical tdTomato^+^ dendrites at DPI-10, presumably some of those contacts were located in the dendritic shaft (Shepherd et al., 2021). However, gephyrin expression in juxtaposition with tdTomato^+^ spines was not obvious until DPI-17, when morphologically distinct spines were first present.

**Figure 4.**
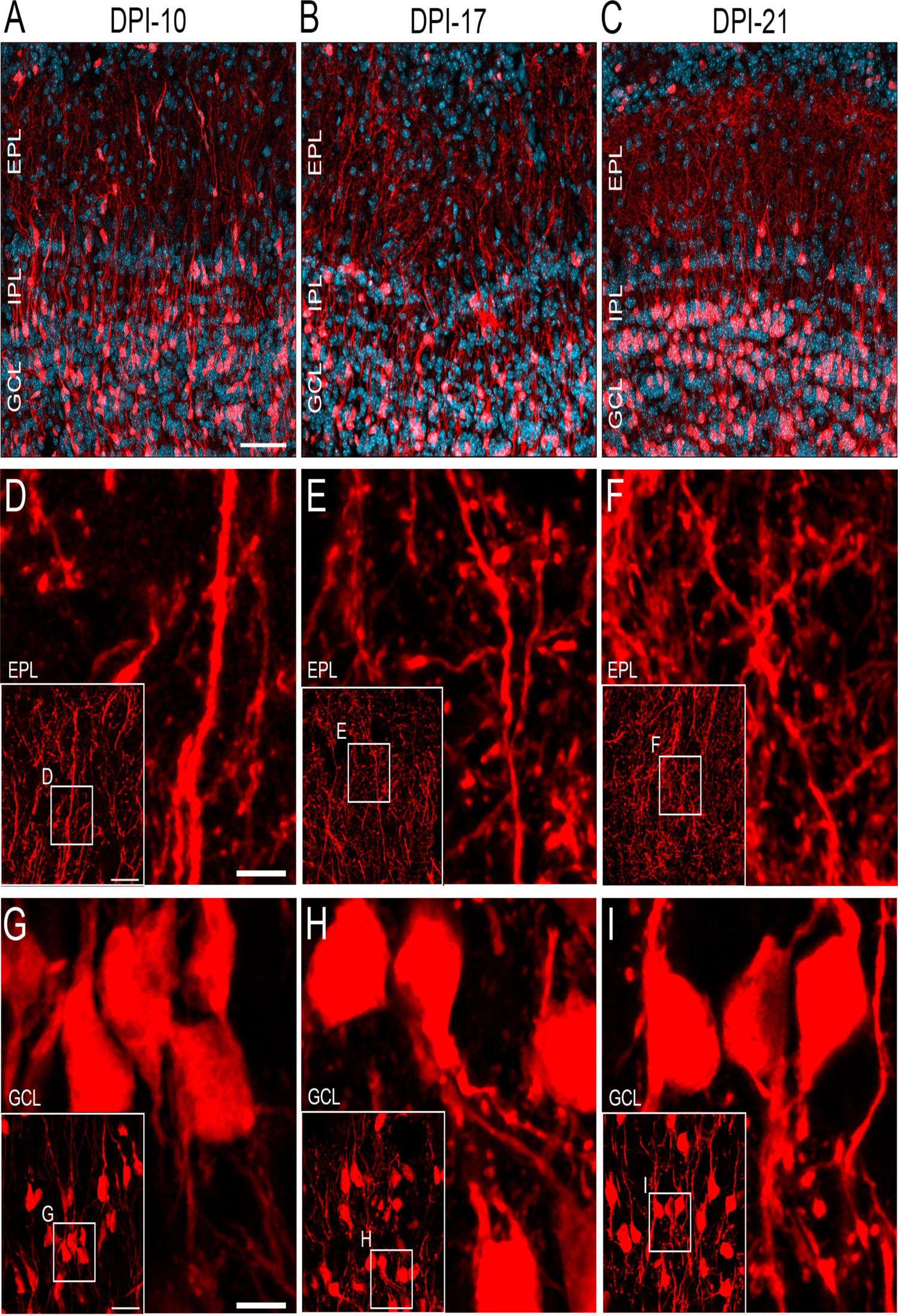
Morphological details of GCs generated at P7. (A-C) Confocal images of OB cross section showing tdTomato^+^ GCs extending their apical dendrites into the EPL 10, 17 and 21 days after 4OH-Tx injection, respectively. Note that at DPI-21 the EPL is highly innervated. (D-F) High magnification confocal images revealing morphological details of GC apical dendrites in the EPL at 10, 17 and 21 DPI. Note the presence of large spine heads and long spine necks at DPI-17 at the earliest. (G-I) Details of obvious GCs basal spiny dendrites at 17 and 21 DPI. Inserts in lower left corner of (D-I) show broader EPL and GCL region from where high magnification pictures were taken. Scale bar (A-C) 50 μm (D-I) 20um. DPI: day post injection, EPL: external plexiform layer, GCL: granular cell layer, IPL: internal plexiform layer.

## DISCUSSION

Our analyses support the hypothesis that timing of neurogenesis represents a key mechanism governing GC fate and integration into neuronal circuits in mice OB. Here, we identified three anatomical configurations of GCs based on when they were generated. These three unique distribution patterns correspond to three distinct developmental moments: 1) embryogenesis/neonatal (P0) stages; 2) P7; and 3) adulthood (>P21). Our data reveal that the distribution pattern of GCs generated at P7 represents a hybrid configuration between those born during embryonic/P0 stages and adulthood. Thus, these data demonstrate that the GCL develops as a continuum following a “outside-in” model from embryonic phases to adulthood. The neurogenic activity continuously provides GCs to the OB that will populate specific sublayers in the superficial-deep axis and integrate into different local circuits depending on timing/age. These data are consistent with and extend prior work stating that adult and perinatally born GC constitute two different functional subpopulations of interneurons involved in different olfactory tasks (Sakamoto et al., 2014a). Additionally, we further characterize those GCs generated at P7, analyzing the timeline of their synaptic integration into the bulbar local circuits. Our results show for the first time that, in the mouse OB, these new interneurons recapitulate the synaptic dynamics of those born in adult stages, receiving glutamatergic inputs first and establishing inhibitory synapses later. However, the tempo at which this sequence of synaptogenesis occurs differs significantly from adult born GCs. Interneurons born at P7 integrate into the immature neuronal circuit earlier than those born into a fully established neuronal network. These findings demonstrate that interneurons produced in different moments of life contribute uniquely to the OB network, with potentially different functional implications, and require significantly shorter time to integrate into the mouse OB circuitry during perinatal phases.

### GCL neurogenesis

Our results lead us to conclude that neurogenesis contributes greatly to the development of the GCL during perinatal phases, while it endows a considerably smaller number of new cells during adulthood (Hinds, 1968; Lemasson et al., 2005). Our data reveal a strong laminar distribution of GCs along the GCL that differs based on timing of neurogenesis. Most embryonic and neonatally generated neurons reside in the MCL and superficial segments of the GCL, whereas the ultimate location of adult born interneurons is preferentially within the deepest portion of the GCL preferentially, and only sparsely observed in the superficial layers. This was unexpected since prior studies have concluded that adult born GCs showed an even distribution from the

MCL to the SEZ or the RMS of the OB (Lemasson et al., 2005). This apparent discrepancy could be explained due to different survival times in both our and Lemasson and colleagues’ studies. Lemasson and colleagues conducted their analysis shortly after administering BrdU in adult animals (20 days), whereas we opted for a significantly longer interval period (50 days). Previous studies showed that 15-45 days is the time window for approximately 50% of adult newborn GCs to undergo apoptosis after developing their dendritic arbors and synaptic sites (Petreanu and Alvarez-Buylla, 2002), and required at least 21 days to establish dendro-dendritic synapses in the EPL (Whitman and Greer, 2007). We suggest that if the survival period were extended beyond 20 days, a considerable proportion of adult newborn cells located in the superficial layers of the GCL, as identified by Lemasson and colleagues, might undergo apoptosis and they would not integrate into the local circuits. This is consistent with previous reports that cell survival rate is largely determined by timing of neurogenesis. Perinatally born neurons may survive and remain in the OB until adulthood, whereas a number of those generated during adult stages are eliminated within few weeks after reaching the OB (Najbauer and Leon, 1995; Biebl et al., 2000; Fiske and Brunjes, 2001; Petreanu and Alvarez-Buylla, 2002; Winner et al., 2002). Our BrdU study revealed that neurons in the GCL originate following a superficial to deep neurogenic gradient, characterized by the “outside-in” pattern, reminiscent of paleocortical structures (Martin-Lopez et al., 2019).

We recognize that a minority of the newborn cells investigated in this study may have been originated locally, as GCs have the ability to be produced within the RMS in the OB (Gritti et al., 2002; Hack et al., 2005; Alonso et al., 2008; Schweyer et al., 2019). However, the potential impact of these locally generated interneurons on the development of the GCL remains uncertain. Earlier research suggested that the local production of GCs may only be noticeable in the initial days after birth, but becomes considerably less significant thereafter (Lemasson et al., 2005). Consequently, the migration and integration into the local circuits of GCs born in the SEZ versus those born in the V-SVZ remain areas requiring further exploration. Nonetheless, the specific origin of these newly produced GCs falls beyond the scope of this study, and we encourage the research community to investigate this aspect further.

As indicated by the results outlined above, the fate of newborn GCs is closely tied to the timing of neurogenesis, suggesting that early-born versus adult-born interneurons may interact with distinct populations of principal cells in the OB. This mirrors previous observations in the somatosensory cortex, where the final position of interneurons is heavily influenced by the timing of neurogenesis (Fairén et al., 1986). Similarly, investigations into cell lineage as a potential determinant of the fate of MGE-derived interneurons have not established a direct link between cell origin and the final location of interneurons in the forebrain (Mayer et al., 2015). Alternate mechanisms, such as the timing of neurogenesis, might have a more significant impact, similar to what has been observed in the case of GABAergic cortical interneurons whose locations are linked to birthdate and phenotype (Rymar and Sadikot, 2007). Further differences between early postnatal and adult-born interneurons have been uncovered. Lemasson and colleagues revealed variations in migration speeds and survival rates of newborn cells in the OB depending on their generation time (Lemasson et al., 2005) and Bugeon et al. (2021) have noted that final location of migrating GCs can be activity-dependent.

Interestingly, we demonstrate an uneven distribution of newborn cells along the GCL superficial-deep axis across ages in both sexes. Sexual dimorphism in the V-SVZ and its impact in neurogenic and apoptotic processes has been studied with controversial results depending on age, strains, techniques employed, etc (Díaz et al., 2009; Kim and Casaccia-Bonnefil, 2009; Oboti et al., 2009; Nunez-Parra et al., 2011; Tatar et al., 2013). However, here we found no evidence of sexual dimorphism in neurogenesis or laminar distribution of GCs generated during embryonic, perinatal, and adult periods within the mouse OB.

Importantly, we identify a cohort of interneurons (P7-born GCs) that cannot be classified as superficial- or deep-GCs based on their hybrid location in the GCL. This makes them an intriguing target of interest to further study their electrophysiological properties and uncover the population of projection neurons with which they interact.

### Integration of newborn GCs into the developing neuronal circuits

While there is an overwhelming literature about adult neurogenesis in the OB, strikingly much less is known about neonatal neurogenesis and the behavior of these early postnatally born interneurons. Here, we show that in the neonatal mouse brain newborn GCs mature and integrate into the local circuits significantly faster than in the adult brain, in accordance with previous studies conducted in our lab (Whitman and Greer, 2007). In the mouse neonatal brain, one study suggested that neuronal activity plays a critical role allowing newborn GCs to reach their final location and ensure their survival (Bugeon et al., 2021). Another study demonstrated that the anatomical organization of progenitor domains within the V-SVZ that give rise to different subpopulations of GCs remains unchanged in both neonatal and adult brains (Merkle et al., 2014). Here, we demonstrate the laminar distribution pattern within the GCL of embryonic, neonatal and early postnatal-generated GCs and show that it differs from one another and also from those observed in the adult mouse brain.

In the rat OB, processes of synaptogenesis and integration into the local circuits differ significantly between perinatal and adult born GCs, which would potentially have different functional implications. In a spatiotemporal analysis of synaptic dynamics, Kelsch and colleagues (Kelsch et al., 2008) uncovered that newborn GCs in both perinatal and adult brain receive glutamatergic inputs before establishing inhibitory synapses from their apical dendrites, as had been previously reported in the adult mouse brain by Whitman and Greer, (2007).

However, the spatial distribution of inputs differs between both ages. New cells in the neonatal brain receive excitatory inputs simultaneously in proximal and distal segments of their apical dendrite. Conversely, in the adult brain excitatory inputs in proximal sections of the apical dendrite precedes synaptic inputs in distal regions of the apical dendrite and basal dendrites (Kelsch et al., 2008).

In the adult mouse OB, the initial synaptic contacts onto newborn GCs occur in their basal dendrites shortly after they depart from the RMS and reach the GCL. At this stage, they receive GABAergic and glutamatergic synapses from axon terminals targeting the GCL (Panzanelli et al., 2009). However, the development of complex dendritic arbors with visible spines in the EPL necessitates a two-week period, and the expression of synaptic markers becomes apparent in the apical dendrites a week later (Whitman and Greer, 2007). To date, there’s been a dearth of data on the synaptic integration timeline of newborn GCs in the early postnatal mouse brain.

Our results reveal for the first time that new GCs in the neonatal OB require less than two weeks to express synaptic markers in the EPL, suggesting that they integrate into the local circuits much faster that those born into the adulthood. Consistent with this, previous studies of neurogenic processes in the dentate gyrus of the hippocampus have shown that the pace of synaptic integration and maturation is notably delayed during adult neurogenesis compared to the speed observed during embryonic and early postnatal neurogenesis (For a review see (Ge et al., 2008).

## Conclusions

The addition of new cells to a developing system entails different challenges compared to integration into a mature adult nervous system. While previous research has focused extensively on adult neurogenesis in the OB (For a review see (Lledo et al., 2006; Lepousez et al., 2013), our efforts here were directed towards understanding the behavior of perinatally born GCs. We conclude that timing of neurogenesis dictates the anatomical configuration of GCs within the GCL, which in turn regulates a preferential synaptic integration with either mitral cell or tufted cell secondary dendrites. Importantly, we uncover that GCs born at P7 exhibit a unique distribution pattern representing a hybrid configuration between those generated during embryogenesis and those produced in the adulthood. Moreover, P7-born GCs integrate into local circuits significantly faster than those born into the adult brain, although they recapitulate synaptic dynamics observed in adult OB neurogenesis. Furthermore, we observe that the distribution pattern of GCs continuously shifts from superficial to deep regions of the GCL over time, with potentially different functional implications. Although beyond the scope of the current study, it would be of interest to study the electrophysiological properties of GCs exhibiting the presented hybrid configuration. Our findings align with previous research indicating that the brain is a plastic organ, continually producing new cells that enable it to adapt to constantly changing external and internal environments. However, the process and tempo at which the newly generated neurons integrate and modify the neuronal circuitry in the OB vary depending on the timing of their birth, potentially resulting in different functional implications.

## Acknowledgements

The authors express their appreciation to Christine Kaliszewski for her creative contributions to processing tissue and data acquisition. The research reported here is supported in part by: The Whitehall Foundation and NIH-NIDCD RO1013791, R01016851 and R01017989 to CAG.

---

**Supplementary Figure 1.**
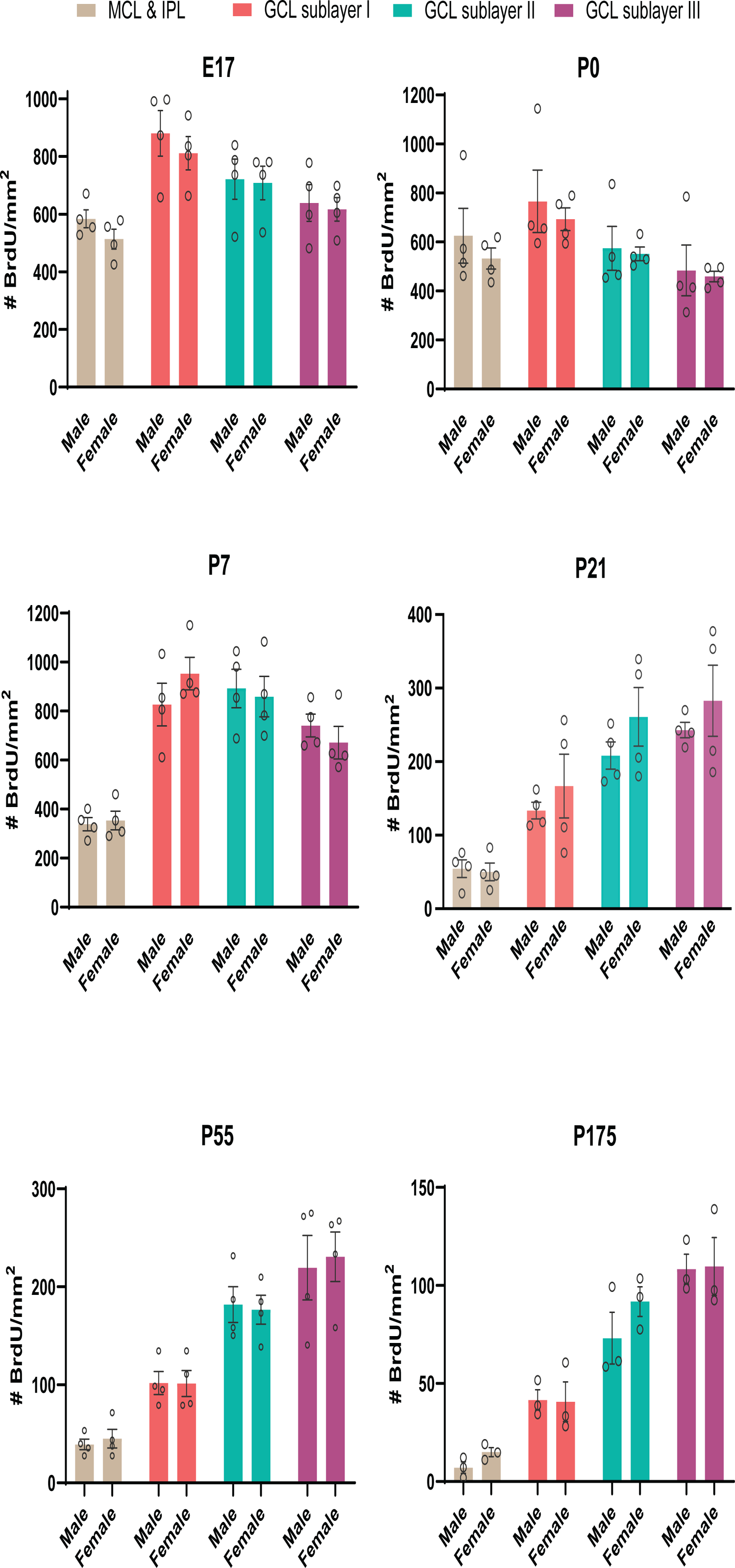
Similar distribution pattern of new GCs produced at six different stages in male and female CD1 mice. Graphics show no significant difference in the number and distribution of newly generated GCs (BrdU^+^) across MCL, IPL and GCL between males and females at any of the timepoint analyzed (E17, P0, P7, P55 and P175).

**Supplementary Figure 2.**
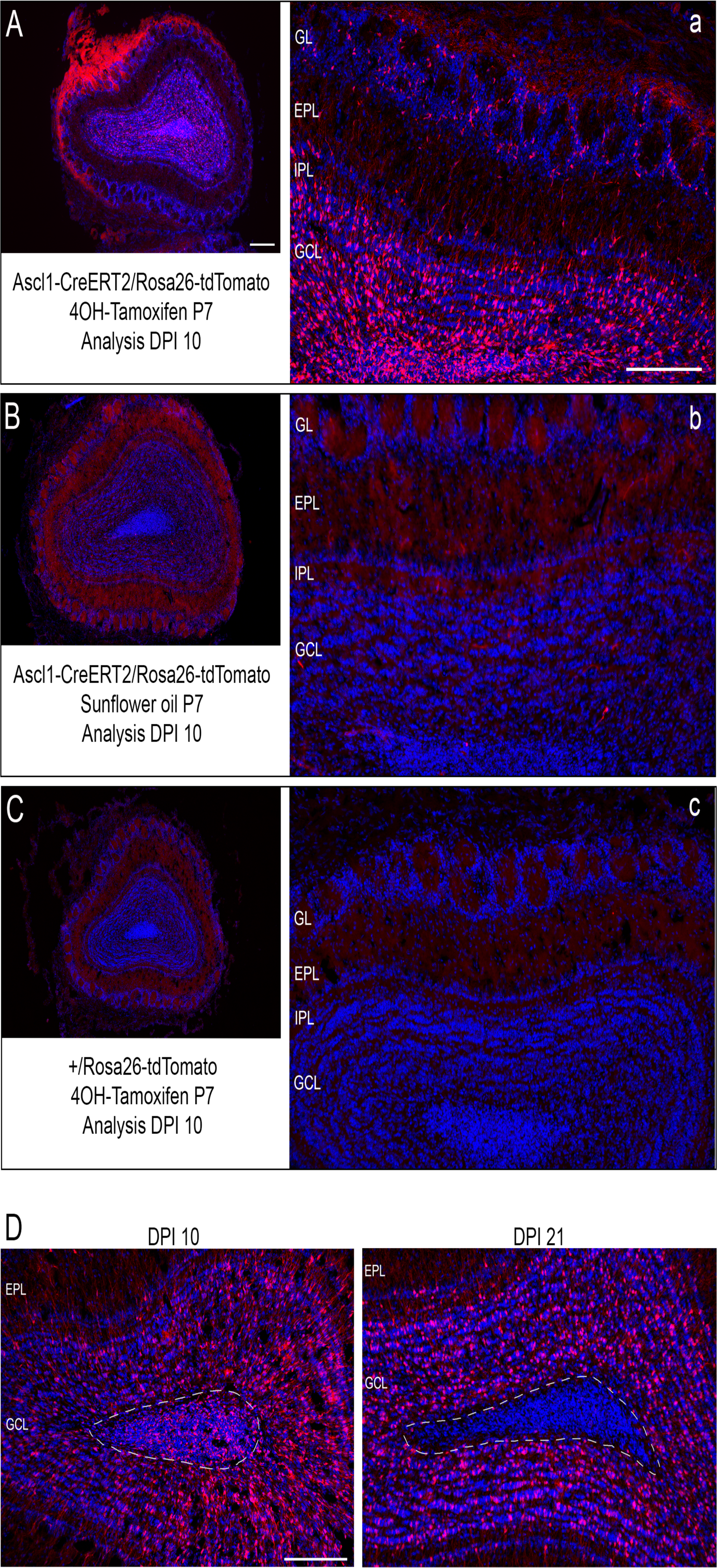
Control experiments for Inducible Cre-lox P System. To test the accuracy of the 4-OH-Tx inducible Cre-LoxP strategy we conducted three control experiments. First, we injected one group of double transgenic mice (Ascl1-CreERT2/Rosa26-tdTomato) with 4OH-Tx at P7 (n = 3) and another group with sunflower oil (vehicle, n = 3). Coronal section of Ascl1-CreERT2/Rosa26-tdTomato mouse injected with 4OH-Tx at P7. Analysis at 10 DPI showed a considerable number of GCs expressing tdTomato (red) in the GCL after 4OH-Tx injection (A), but not after sunflower oil injection (B). Likewise, double transgenic animals injected with 4OH-Tx showed tdTomato^+^ apical dendrites extended into the EPL (Figure 3. a), unlike those injected with sunflower oil (B). We also observed another group of tdTomato^+^ interneurons located in the GL (A and a), which were not included in our study. Finally, a third group of reporter mice with +/Rosa26-tdTomato genotype was injected with 4OH-Tx (n = 3). Ten days after 4OH-Tx injection no tdTomato^+^ GCs were observed in the GCL in the absence of CreERT2 (C and c). To test that tdTomato is transiently expressed in OB neuronal precursors in the presence of 4OH-Tx, we analyzed OB coronal sections of double transgenic mice injected at P7 with 4OH-Tx 10 and 21 DPI. We observed tdTomato^+^ cells in the subependymal zone 10 DPI, but no 21 DPI (D) corroborating that the expression of tdTomato is transient and specific for OB neuronal precursor cells that give rise to interneurons in the mouse postnatal brain. Therefore, this in vivo fate tracing strategy allows us to specifically study GCs born specifically during a very narrow time window. Scale bars: (A-D) 250 μm, (a-c) 200 μm.

